# Scalable estimation of microbial co-occurrence networks with Variational Autoencoders

**DOI:** 10.1101/2021.11.09.467939

**Authors:** James T. Morton, Justin Silverman, Gleb Tikhonov, Harri Lähdesmäki, Rich Bonneau

## Abstract

Estimating microbe-microbe interactions is critical for understanding the ecological laws governing microbial communities. Rapidly decreasing sequencing costs have promised new opportunities to estimate microbe-microbe interactions across thousands of uncultured, unknown microbes. However, typical microbiome datasets are very high dimensional and accurate estimation of microbial correlations requires tens of thousands of samples, exceeding the computational capabilities of existing methodologies. Furthermore, the vast majority of microbiome studies collect compositional metagenomics data which enforces a negative bias when computing microbe-microbe correlations. The Multinomial Logistic Normal (MLN) distribution has been shown to be effective at inferring microbe-microbe correlations, however scalable Bayesian inference of these distributions has remained elusive. Here, we show that carefully constructed Variational Autoencoders (VAEs) augmented with the Isometric Log-ratio (ILR) transform can estimate low-rank MLN distributions thousands of times faster than existing methods. These VAEs can be trained on tens of thousands of samples, enabling co-occurrence inference across tens of thousands of microbes without regularization. The latent embedding distances computed from these VAEs are competitive with existing beta-diversity methods across a variety of mouse and human microbiome classification and regression tasks, with notable improvements on longitudinal studies.

## 1 Introduction

Understanding microbe-microbe interactions is one of the major outstanding questions in microbial ecology. Microbes are suspected to interact through ecological relationships, whether it be through the exchange of metabolic resources through cross feeding, through competition over resources or through neutral interactions. Furthering our knowledge of microbe-microbe interactions has broad implications for designing and perturbing microbial communities for bioengineering [1], drug design and clinical trial applications [2]. Inferring these interactions is complicated by the unknown complexity behind microbial metabolism and consequentially the vast number of microbes that have not yet been cultured. Multiple studies have constructed microbial co-cultures [3, 4], but the logistical challenge of setting up these approaches limits the number of co-culture assays that can be conducted, which pales in comparison to the combinational number of potential microbe-microbe interactions.

High throughput sequencing has presented an alternative means to understand microbe-microbe interactions. Amplicon sequencing and shotgun metagenomics sequencing present the possibility of counting microbial individuals within a biological specimen, and recent efforts have catalogued and assembled genomes across millions of unculturable microbes. With these measurements, it is possible to perform correlation analysis to narrow down potential microbe-microbe interactions to validate for co-culturing [3]. However, metagenomic datasets present numerous statistical challenges that hamper inference and interpretation, including high dimensionality, high sparsity and compositionality where units of concentrations or cells per gram are often not collected. The lack of scale information is particularly problematic; since only microbial proportions can be estimated, the collected data are confined under a simplicial geometry. This projection limits the conclusions that can be drawn regarding microbial interactions, since the simplex imposes a negative bias on microbe-microbe correlations. This promotes excess false positives and false negatives when applying conventional correlation methods [5, 6, 7].

Over the last decade, there have been numerous attempts to develop statistical estimators of microbe-microbe correlations. SparCC [7] proposed to infer absolute correlations by exchanging the scale identifiability issue with a heuristic sparse correlation assumption. SPEIC-EASI [8] and later gCODA [9], MP-Lasso [10] and Flashweave [11] proposed to solidify the sparse correlation assumption with sparsity-inducing graph regularization. However, since compositional approaches are designed for dense data, these methods require an imputation strategy, which is problematic when dealing with very sparse data. More recent works have proposed to utilize the Logistic-Normal distribution in combination with a counting distribution such as the Multinomial or the Poisson to account for the sparsity issue [12, 13, 14, 15, 16, 17, 18, 19, 20, 21]. However, due the non-conjugacy between counting distributions and the Normal distribution, Bayesian inference based on Markov Chain Monte Carlo (MCMC) posterior sampling is challenging to scale-up for high dimensional microbiome datasets. Variational approximations have been developed to overcome these computational issues [15, 20, 21], utilizing normal approximations or quadrature methods to handle the conditional non-conjugacy between these count distributions and the logistic normal distribution. While these approaches are much faster than MCMC, they may introduce additional estimation bias [22]. Furthermore, none of these methods have been applied to datasets with more than a thousand samples, requiring either excessive filtering of microbes or regularization on the logistic-normal covariance matrix to avoid identifiability issues in the low sample size regime.

In the machine learning community, Variational Autoencoders (VAEs) have been proposed as a scalable means around these non-conjugacy issues. By using reparameterization gradients [23], it is possible to estimate the posterior distribution of the parameters of interest by directly evaluating and maximizing the Evidence Lower Bound (ELBO) with stochastic gradient descent. More recently, connections between VAEs and Probabilistic PCA (PPCA) have been identified [24], where Linear VAEs can estimate the same principal components as PPCA. However, the connection between VAEs and PPCA is currently limited to Gaussian distributed data and not well-suited for count data. Showing that VAEs can recover the correct principal components from count data is nontrivial due to the non-conjugacy issues between the logistic normal distribution and count distributions such as the multinomial distribution. Furthermore, the parameters of the multinomial distribution are compositional; they are constrained within the simplex and the resulting covariance matrix is singular and non-invertible [6, 25]. Aitchison provided a means to perform PCA on compositional data [26] and more recent efforts showed how to obtain principal components through unconstrained inference using the Isometric Log-ratio (ILR) transform [27, 28].

### Our contribution

Here we show that VAEs can be modified to estimate Multinomial Logistic Normal (MLN) distributions with low-rank covariance matrices. Rather than utilizing coordinate ascent variational inference as shown in Poisson PCA [15], we directly sample the ELBO with reparameterization gradients and show that this approach can scale to datasets with tens of thousands of samples and ten thousand microbial species. We make the case that the ILR transform is critical for estimating the principal components in multinomial logistic normally distributed data. This provides an opportunity to utilize compositional data analysis techniques to interpret the resulting embeddings and extract microbial co-occurrence information.

## 2 Methods

### 2.1 The connection between Linear VAEs and Probabilistic PCA

VAEs were originally proposed as a generative model [23], and are now commonly deployed across scientific disciplines, making contributions to single-cell RNA sequencing [29], microbiome modeling [30], protein modeling [31, 32, 33], natural language processing [34] and image processing [23]. Lucas et al. [24] has previously shown that the following two models can obtain the same maximum likelihood estimates of principal components ***W*** :

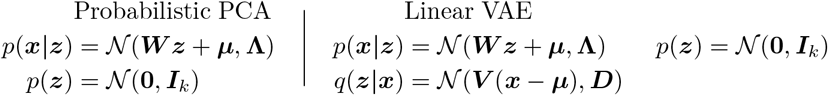

Here, ***x*** ∈ ℝ^*d*^ represents the real-valued *d* dimensional observations, ***z*** ∈ ℝ^*k*^ represents the corresponding latent representation reduced down to *k* latent dimensions. *p*(***x***|***z***) denotes the likelihood of observations ***x*** given the latent representation ***z*** ∈ ℝ^*k*^, *p*(***z***) denotes the prior on ***z***. In Tipping et al, **Λ** = *σ*^2^***I*** for *σ*∈ ℝ, and parameters of interest, namely the latent representation ***z*** and the principal components ***W*** ∈ ℝ^*d×k*^, are directly estimated through Expectation-maximization [35].

It was shown that Linear VAEs can directly predict the posterior of ***z*** given by *q*(***z***|***x***) with a neural network rather than than directly estimating ***z*** as additional learnable parameters [24]. As a result, the number of model parameters does not scale with the size of the input; instead only a projection matrix ***V*** ∈ *ℝ*^*k×d*^ and a diagonal covariance matrix ***D*** ∈ ℝ^*k×k*^ are needed approximate *q*(***z***|***x***). The parameters underlying Linear VAEs are estimated by maximizing the ELBO

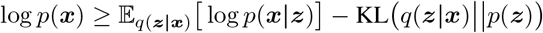

For Linear VAEs with a Gaussian likelihood, the variational posterior distribution *q*(***z***|***x***) can be shown to analytically agree with the posterior distribution *p*(***z***|***x***) learned from PPCA [24]. However, deriving this connection for count-based likelihoods such as the multinomial distribution is complicated due to non-conjugacy issues (Appendix A.2). This is further complicated by a missing definition of Multinomial PPCA.

### 2.2 Multinomial Probabilistic PCA

One challenge of defining Multinomial PPCA is defining a bijection between proportions and logits that leads to scale identifiability. The “softmax-trick” highlights such a identifiability issue where any specified shift constant ***a*** = (*a*, · · ·,*a*) ℝ^*d*^ applied to logits ***x*** ∈ ℝ^*d*^ will not change the output of softmax *ϕ*(***x***), namely *ϕ*(***x***) = *ϕ*(***x*** + ***a***). As highlighted in the compositional data analysis literature [26, 28], softmax is a degenerate function and not accounting for this identifiability issue will confound principal components estimation (i.e. *ϕ*needs to be isomorphic). Another challenge is the estimated loadings needs to encode the covariance matrix across dimensions, meaning that *ϕ*needs to preserve row distances of ***x*** (i.e. *ϕ*needs to be isometric).

The inverse Isometric Log-ratio (ILR) transform [27] is such a function that satisfies both isomorphism and isometry when transforming logits in ℝ^*d−*1^ into proportions in the simplex 𝕊^*d*^. Due its attractive properties, the ILR transform has been shown to be more suitable for principal components analysis [28]. The ILR and inverse ILR are given as follows:

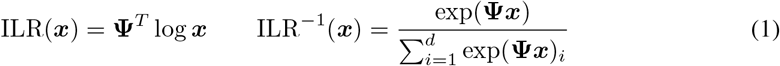

where **Ψ** ∈ ℝ^*d×d*−1^ is a basis such that **Ψ**^*T*^ **Ψ** = ***I***_*d* − 1_ and 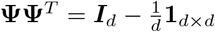. Any orthonormal basis can parameterize the ILR transform and some of these bases can be represented by binary trees [36, 37, 38]. For our model, we use a phylogenetic tree, due to its biological interpretability and low runtime requirements. See Appendix A.1 for further discussions on the ILR.

Letting *ϕ*(***x***) denote the inverse ILR transform, PPCA can then be extended to multinomially distributed data with the following generative model:

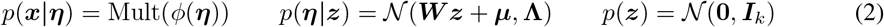

Here ***W*** ∈ ℝ^*d−*1*×k*^ represents the principal components, and **Λ** ∈ ℝ^*d−*1*×d−*1^ is a diagonal covariance matrix. For a single sample, ***x*** ∈ ℕ^*d*^ are the observed *d* -dimensional counts, ***η*** ∈ ℝ^*d−*1^ are the latent logits and ***z*** ∈ ℝ^*k*^ is the latent representation. Unlike standard PPCA, the principal components ***W*** cannot be used project heldout data to the latent space since the ILR transform cannot be directly applied to counts (i.e. log(0) is not defined). Furthermore, the estimation of the Multinomial PPCA parameters is nontrivial. The term *ϕ*(***η***) follows logistic normal distribution *ϕ*(***η***) ∼ ℒ 𝒩 (***W z*** +***μ*, Λ**), as shown by Aitchison [39], as a result the expectations of *ϕ*(***η***) are not analytically tractable, complicating the application of Expectation-maximization. Furthermore, *p*(***x***|***z***) yields a MLN distribution, which is given by marginalizing out ***η*** in *p*(***x***|***z***) = ∫_***η***_ *p*(***x***|***η***)*p*(***η***|***z***)*d****η***. This integral is not tractable; as a result, this distribution does not have an analytically defined probability density function, complicating maximum likelihood estimation. There have been multiple attempts to estimate the posterior distribution with MCMC [40, 17, 14, 12], but the complexity of this distribution requires a large number of Monte Carlo samples, limiting the scalability of these methods. Other efforts have developed variational methods have utilized loose lower bounds of the ELBO to combat the conditional non-conjugacy [41, 20, 21]. However these approximations have been shown to be suboptimal to their Monte Carlo counterparts [22].

### 2.3 Multinomial Variational Autoencoders

Here, we argue that a Multinomial VAE can address these non-conjugacy issues by using reparameterization gradients. The concept of a Multinomial VAE have been previously proposed in the context of recommender systems [42]. However, these architectures contain the scale identifiability issue discussed above, preventing the estimation of principal components and the resulting covariance matrix. As suggested earlier, the scaling identiability issue between the logits ***η*** and the multinomial proportions can be resolved with the inverse ILR transform with the following reformulation:

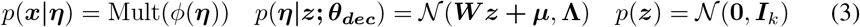

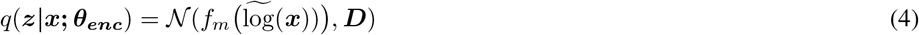

where ***θ***_***dec***_ = {***W***, **Λ**} denotes the decoder parameters, ***θ***_***enc***_ = {*f*_*m*_, ***D****}* denotes the encoder parameters and ***μ*** ∈ ℝ^*d−*1^ is a bias parameter. We relax the definition of **Λ** to be any diagonal covariance matrix in order to provide more flexibility in modeling overdispersion. Here, *q*(***z***|***x*; *θ***_***enc***_) denotes the variational posterior distribution of ***z*** given by the encoder *f*_*m*_ represented as an *m*-layer dense neural network with appropriate activations. This encoder is directly used to evaluate *p*(***η***|***z*; *θ***_***dec***_). Furthermore, flat priors are assumed for all variables except ***z***. The encoder architecture is memory efficient; by approximating the posterior distribution of ***z***, the encoder provides a way to project counts into the latent space without memorizing ***z***. Unlike like previous MLN estimators [12, 16, 17], the number of learnable parameters is fixed with respect to the number of samples. If a single layer encoder is defined, then maximizing the ELBO becomes a biconvex optimization (Appendix A.4). Furthermore, simulations suggest that the Multinomial VAEs exactly recovers the Multinomial PPCA parameters when applied to completely dense data.

Estimating principal components becomes more problematic when dealing with sparse count data, since logarithms are not defined for zeros. A common approach to this problem is to introduce a pseudocount before applying a logarithm. To ensure scale invariance, we normalize the inputs to proportions after adding a pseudocount, namely 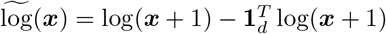 The choice of pseudocount is arbitrary and can introduce unwanted biases. To alleviate this issue, we introduce a nonlinear, possibly deep encoder neural network *f*_*m*_ highlighted in Equation 4; we expect that the universal approximation theorem would apply here [43, 44] where the representation quality of ***z*** will improve with more complex neural networks. This is supported in our simulation benchmarks; more complex encoder architectures can lead to better estimates of principal components on sparse data.

## 3 Results

To showcase the merits of Multinomial VAEs, we showcase its performance across a wide variety of benchmarks. In Section 3.1 we showcase scenarios where Multinomial VAEs can recover Multinomial PPCA parameters. In Section 3.2 we demonstrate that Multinomial VAEs are competitive to state-of-the-art beta-diversity metrics in the context of a mouse fecal pellet dataset with 11k samples, 5k microbes and on a human fecal sample dataset with 25k samples, 11k microbes. In Section 3.3 we extract the embedding information from the Multinomial VAEs to demonstrate that it can learn meaningful ecological relationships.

### 3.1 Multinomial VAEs recovers Multinomial PPCA loadings in simulations

To investigate the connections between Multinomial VAEs and Multinomial PPCA, we constructed multiple simulations using the Multinomial PPCA generative model in Equation 2. The agreement between the estimated and ground truth principal components was measured with two metrics, namely pairwise Pearson correlation between the estimated and ground truth correlation matrices and axis alignment metric between the ground truth and estimated principal components.

The correlation matrices are estimated through the inner product of the VAE decoder weights, namely ***W W***^*T*^. The axis-alignment metric is given by the average cosine distance between estimated and ground truth eigenvectors. If the data has no zeros, the Multinomial VAEs can exactly recover the ground truth correlations generated from Multinomial PPCA (Figure 1a). However, as more sparsity is induced, it becomes more difficult to estimate the ground truth correlations (Figure 1b and d). However, having a deeper encoder can lead to higher fidelity principal components (Figure 1d), implying that higher complexity encoders may be needed to estimate the latent embeddings ***z*** for sparse datasets.

**Figure 1:**
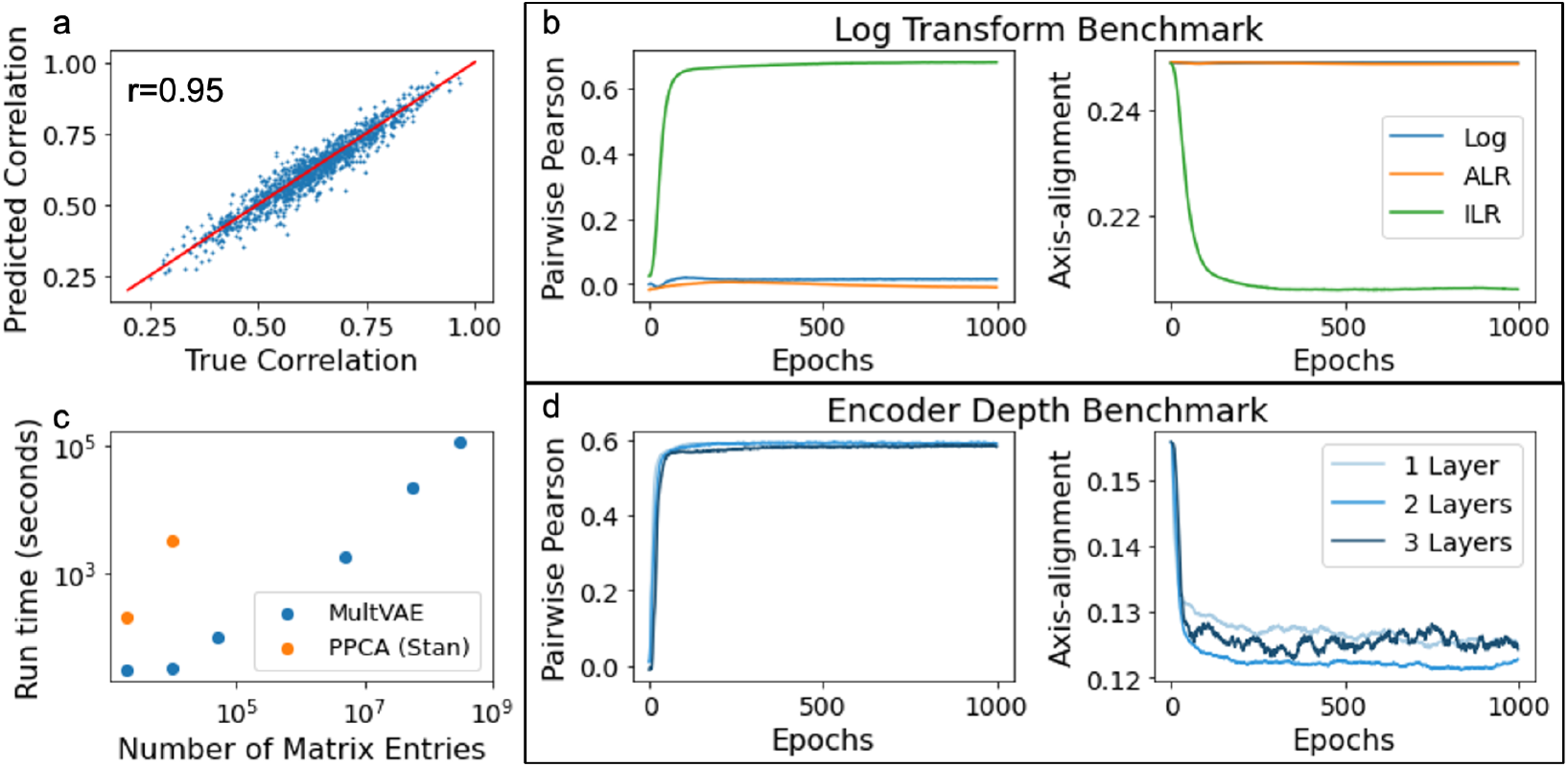
Multinomial PCA simulation benchmarks: (a) Multinomial VAEs can recover the ground truth principal components and correlations if there are no zero counts present in the dataset. (b) Multinomial VAEs augmented with the ILR transform outperforms the ALR and Log transforms on a simulation dataset with 40% sparsity and a single layer encoder. (c) Multinomial VAEs are orders of magnitude faster than Multinomial PPCA fitted with Hamiltonian Monte Carlo [45]. (d) Another simulation showing that Multinomial VAEs trained with multiple encoders can obtain higher fidelity principal components. Lower axis-alignment metric indicates more agreement between the estimated and ground truth principal components with respect to their directionality.

We also find that the ILR transform plays a critical role in recovering the ground truth principal components. The ILR transform obtains significantly better estimates of the ground truth correlations compared to VAEs using the additive log-ratio (ALR) transform and the standard Log transform (Figure 1b). This observation is consistent with the compositional data analysis literature; the ALR transform and its corresponding inverse do not preserve isometry and consequentially does not recover the ground truth correlations [46]. Most VAEs constructed to analyze single-cell RNAseq data, including scVI [29] and LDVAE [47], use the standard softmax function to convert logits to proportions. Our simulation benchmarks suggest that this is a suboptimal design choice if one wishes to recover the underlying correlations from sequencing count data using a linear decoder.

One major benefit of using Multinomial VAEs over previous MLN estimators is the speed of training. We attribute this speed up due to the reparameterization trick. Due to the ability to deploy stochastic gradient descent, the non-conjugacy between the Multinomial and Logistic normal are no longer the bottleneck in training. Furthermore, the utilizing the GPU with Pytorch [48] provides an additional speedup. Even though it can take thousands of epochs to train the Multinomial VAE [49], the Multinomial VAE can handle datasets orders of magnitude larger than what Multinomial PPCA can be fit to by using state-of-the-art MCMC software [45] (Figure 1c).

### 3.2 Multinomial VAEs are competitive in distinguishing mouse and human phenotypes

We trained the Multinomial VAE on a mouse dataset and a human dataset that were uploaded to Qiita [50]. Our pretrained Multinomial VAEs were then compared against state-of-the-art beta-diversity methods, including Bray-Curtis and Unifrac [51] in addition to the Linear VAE implemented in scVI (LDVAE) [29, 47] and Latent Dirichlet Allocation (LDA) [52] on the mouse dataset. For the human dataset, only the Multinomial VAE, Bray-Curtis and Unifrac were benchmarked.

Since all of the above methods are unsupervised, K nearest neighbors classifiers were trained to distinguish mouse phenotypes and human phenotypes on the held out samples. Across all of the classification tasks, the Multinomial VAEs obtains competitive classification accuracy, with a significant improvement on two longitudinal datasets [53, 54] and in age prediction [59] (Figure 2). Similar conclusions were drawn with compositional PCA [60] and compositional tensor factorization [61], where robust composition metrics were able to distinguish longitudinal samples more easily than non-compositional metrics. As hinted by the simulation results, we suspect that this performance difference can be explained by the lower sparsity in these longitudinal datasets. On the other hand, we see the Multinomial VAE sometimes underperforms Bray-Curtis and Unifrac when distinguishing IBD phenotypes.

**Figure 2:**
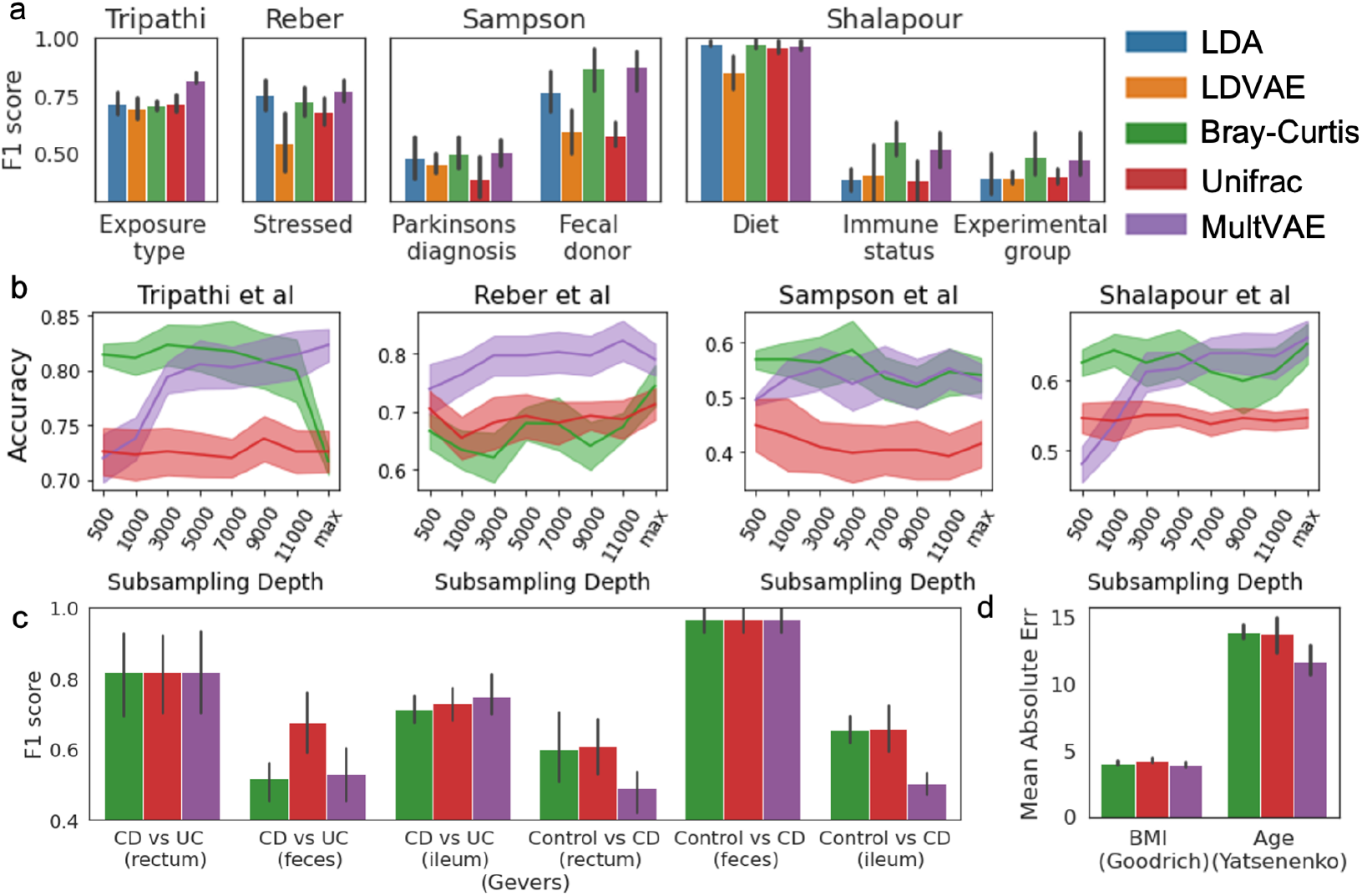
Classification benchmarks : (a) Classification across multiple tasks with LDA, LDVAE, Bray-Curtis, Unifrac and MultVAE across 4 mouse datasets [53, 54, 55, 56]. (b) Sequencing depth benchmarks with Bray-Curtis, Unifrac and MultVAEs. (c) Classification benchmarks on the Gevers et al dataset [57]. (d) Regression benchmarks to predict body mass index (BMI) in the Goodrich et al dataset [58] and age in the Yatseneko et al dataset [59]

We also saw that LDVAE under-performs all of the methods benchmarked here. While it is plausible that the standard softmax could be contributing to its under-performance, we suspect that there are other design choices that further degrades performance. LDVAE uses the encoder to estimate both feature-specific overdispersion parameters and sample-specific variances, both of which induces an identifiability issue between the encoder and the decoder leading degraded representation quality [24]. Furthermore, the zero-inflated negative binomial may not an appropriate distribution for sparse microbiome datasets, since it will lead to abundance overestimation [62].

To understand the Multinomial VAE performance, we investigated how the classification accuracy improves as the sequencing depth is increased. The question of normalization has been a controversial topic in the microbiome literature [63]. One of the commonly used normalization techniques is rarefaction, where sequencing counts across samples are subsampled to the same sequencing depth. This approach is problematic since it is stochastic and increases uncertainty, often throwing out more than 90% of reads, but has been shown to reduce the confounding variation due to sequencing depth variation [64]. As hinted by our benchmarks, Bray-Curtis and Unifrac classification accuracy does not improve as more sequencing reads are added, whereas the Multinomial VAE classification accuracy strictly improves with higher sequencing depth. We attribute this difference due to the Multinomial distribution’s ability to compute sequencing depth uncertainty. Only the Multinomial VAE incorporates counting uncertainty into the model, which is highly desirable for handling samples with varying sequencing depths.

Aside from normalization, one of the issues when performing meta-analysis on multiple datasets is the presence of batch effects, where unobserved confounders complicates the comparison of multiple datasets. Technical variation due to sampling and processing protocols is one of the sources of batch effects. Whereas the studies in the mouse dataset used consistent protocols, there were some differing sample storage protocols in the human dataset. To justify training the Multinomial VAE on the mouse and the human datasets, we benchmarked the pretrained Multinomial VAE on technical replicates across multiple protocols (Table 1) and compared it against Combat [69] and Combat-seq [70]. While the Multinomial VAE itself is not a batch correction method, we found that it performed comparably to Combat and Combat-seq. Although the Multinomial VAE displayed noticeable batch effects in the MBQC dataset, the effect is significantly outweighed by the differences across biological samples. Furthermore, the Multinomial VAE was able to distinguish the different biological samples in the MBQC dataset comparable to Combat and even better than Combat-seq. Previous literature showed noticeable batch effects due to sample collection method [71], but we found that if we focus on samples that were either immediately refrigerated or flash frozen or stored with RNAlater, the storage effects were significantly smaller than the biological variation across batch correction methods across all three methods.

**Table 1:**
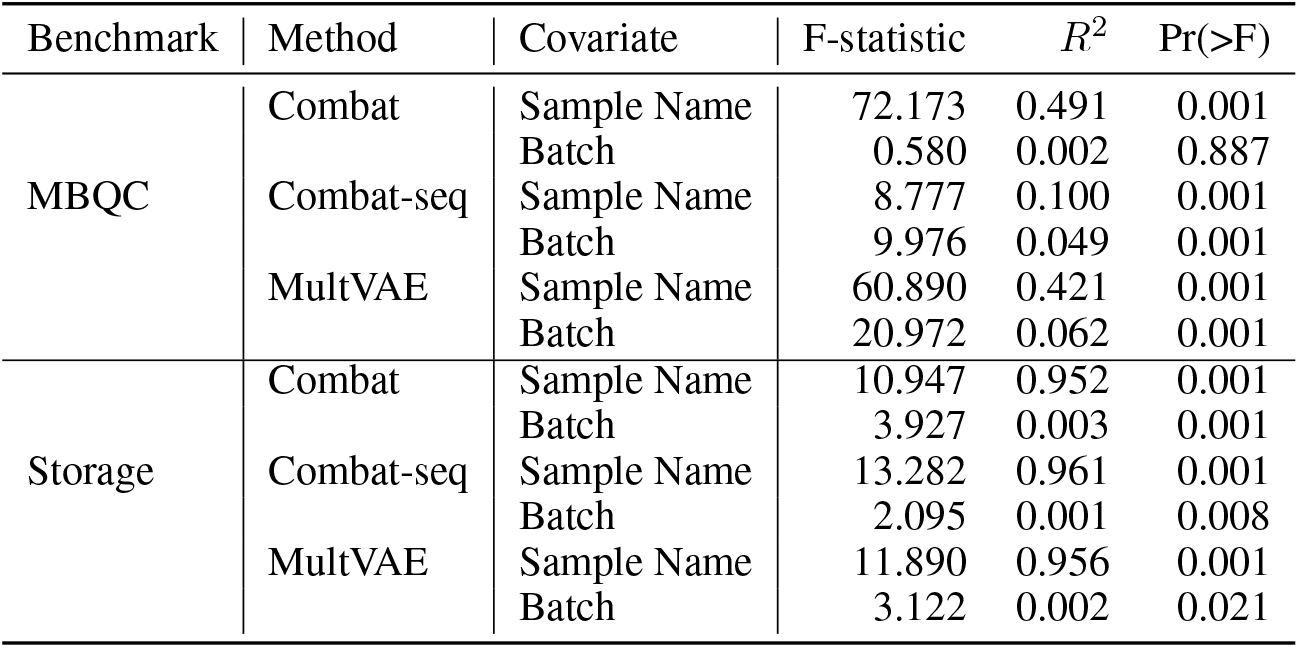
Batch effects benchmark in the MBQC [65] and the Mayo storage studies [66, 67]. Multinomial VAEs, Combat and Combat-seq evaluated to differentiate the biological samples and the technical effect of processing lab and storage conditions with Permanova [68]

| Benchmark | Method | Covariate | F-statistic | $R^2$ | Pr(>F) |
| --- | --- | --- | --- | --- | --- |
| MBQC | Combat | Sample Name | 72.173 | 0.491 | 0.001 |
|  |  | Batch | 0.580 | 0.002 | 0.887 |
|  | Combat-seq | Sample Name | 8.777 | 0.100 | 0.001 |
|  |  | Batch | 9.976 | 0.049 | 0.001 |
|  | MultVAE | Sample Name | 60.890 | 0.421 | 0.001 |
|  |  | Batch | 20.972 | 0.062 | 0.001 |
| Storage | Combat | Sample Name | 10.947 | 0.952 | 0.001 |
|  |  | Batch | 3.927 | 0.003 | 0.001 |
|  | Combat-seq | Sample Name | 13.282 | 0.961 | 0.001 |
|  |  | Batch | 2.095 | 0.001 | 0.008 |
|  | MultVAE | Sample Name | 11.890 | 0.956 | 0.001 |
|  |  | Batch | 3.122 | 0.002 | 0.021 |

### 3.3 Multinomial VAEs recover ecologically meaningful relationships

To further motivate the architecture underlying Multinomial VAEs, we argue that there is a connection between Multinomial VAEs and compositional PCA [26]. Because the logits ***η*** characterizing the multinomial parameters are embedded in ILR space, by design the decoder weights ***W*** are expected to hold similar interpretation to the loadings estimated in compositional PCA. Namely, we expect

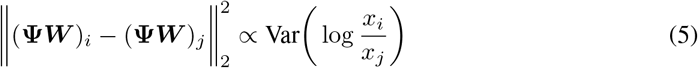

where ***x*** refers to the observed proportions and *i* and *j* refer to the two features being compared. The variance log ratio (VLR) [72, 73] holds a simple interpretation: if the VLR is close to zero, that means the two microbes are tightly co-occurring. Furthermore, large VLR values indicate microbes that aren’t often observed together in the same biological samples. For highly abundant microbes, we see there is a proportional relationship between VLR and the VAE decoder distances (Figure 3), suggesting that Multinomial VAEs do indeed estimate microbial co-occurrences. Empirically, we do observe that there isn’t an exact agreement between the VAE decoder distances and the VLR. This could be explained by the differences between the two methodologies; Multinomial PPCA accounts for count uncertainty and zeros whereas compositional PCA cannot handle zeros.

**Figure 3:**
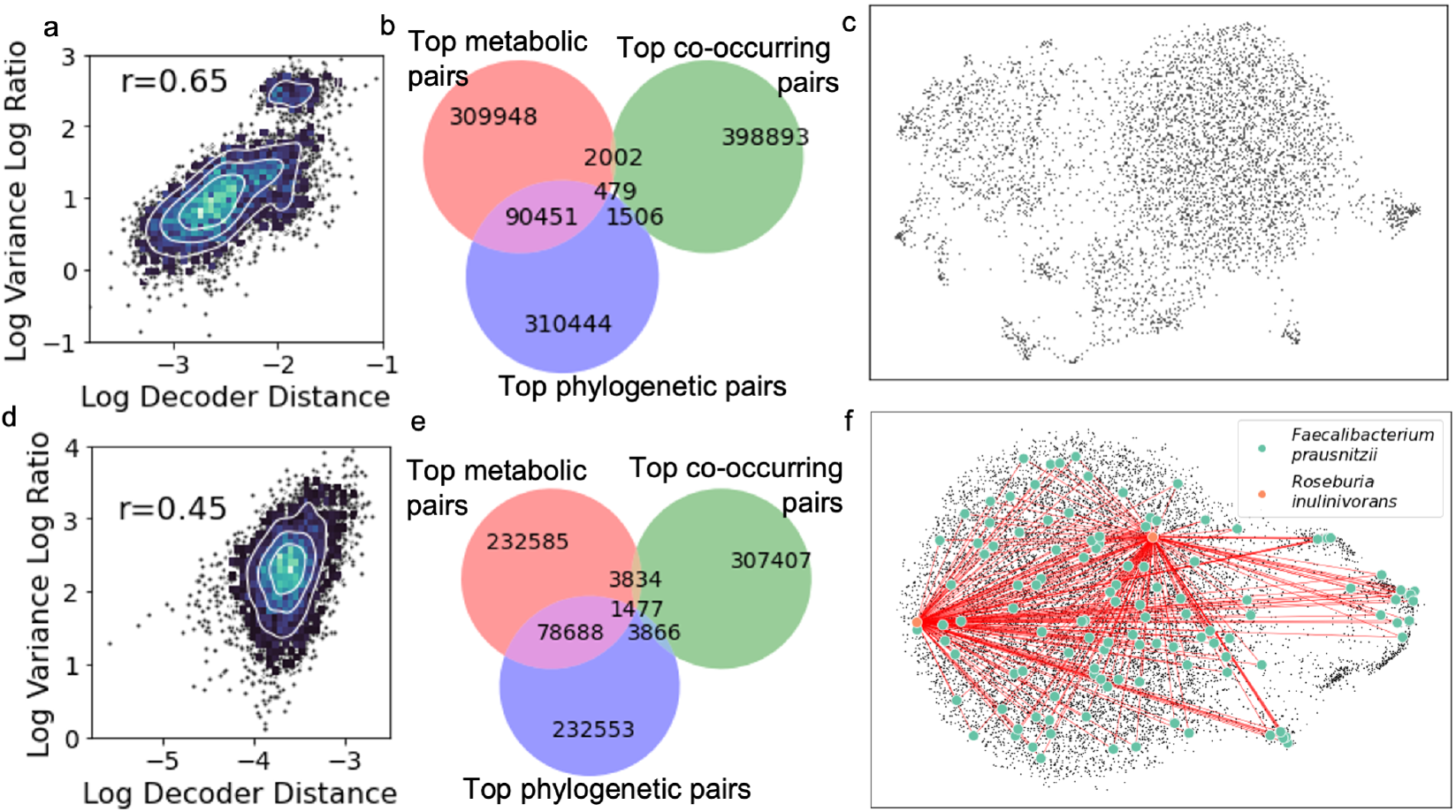
Microbial co-occurrences extracted from mouse and human datasets. (a, d) Euclidean distances extracted from VAE decoder is proportional to the variance log ratio metric across the top 100 most abundant microbes. (b, e) Venn diagrams looking at top *K* microbial pairs according to pathway similarity, phylogenetic similarity and co-occurrence probability. *K* is determined by the number of pairs of taxa that have identical KEGG profiles. (c, f) UMAP [74] plot of all of the microbes embedded using cosine distance with co-cultured microbes highlighted. Cosine distance was used instead of Euclidean distance due to ease of interpretation [75]; a cosine distance of 0 indicates highly co-occurring microbes, 1 indicates neutral interactions and 2 indicates highly anti-co-occurring microbes. The UMAP embeddings for the human gut microbiome interactions are overlapped with *Faecalibacterium prausnitzii* and *Roseburia inulinivorans* putative interactions that was experimentally validated in Das et al [3].

To identify potential driving patterns underlying these interactions, we extracted pathway information via Picrust2 [76] in addition to computing phylogenetic distances between microbes. We found pairs of microbes that have complete overlap in KEGGs only account for a fourth of those microbes were amongst the top phylogenetically similar microbes. We suspect that this hints at the annotation incompleteness of KEGG [77], where pathways from under-studied organisms are under-represented. Interestingly we found that microbes that had similar pathways and microbes that are phylogenetically similar are not amongst the top co-occurring microbes. This observation is consistent with Darwin’s phylogenetically limiting hypothesis [78]; microbes that are phylogenetic similar tend to have similar enzyme profiles, putting them in direct competition since they consume similar resources.

This pattern is also observed in both *Faecalibacterium prausnitzii* and *Roseburia inulinivorans*. These two microbes have been observed to exhibit a cross-feeding relationship in co-culture experiments [3], and also exhibits wide reaching strain diversity. *Faecalibacterium prausnitzii* in particular has dozens of strains detected and these strains are also spread across the k-nearest neighbors graph. This provides further evidence that phylogenetically similar microbes tend not to co-occur. In fact, our analysis suggests that almost no microbes were detected to be tightly co-occurring; none of the microbial pairs had a cosine distance outside of [0.8, 1.2] (Figure S2). The reasons underlying this observation are currently not clear to us; it is plausible that because most of the studies are cross-sectional, mostly neutral interactions are observed [79] where the gut ecosystems act largely independent of each other. It is important to note that the spatial-temporal nature of these interactions are not measured in this analysis, and as a result the conclusions that can be drawn from these estimated microbe-microbe correlations are limited. Obtaining spatial resolution on the colon [80] or dense time series sampling could help with improving microbial correlation estimates. We anticipate that the Multinomial VAEs presented here can serve as a scaffold for such future efforts.

## 4 Discussion

Most inquiries about the statistical properties surrounding microbiome data have been limited to small sample sizes. Due to the high dimensionality of these datasets, feasible inference required assumptions such as sparsity or imputation that are not easily verifiable. By assembling a large collection of datasets, this is the first study of its kind to bring these types of assumptions to question. Our estimated decoder weights did not follow a sparse distribution (Figure S2), bringing into question the merits of sparsity inducing distributions like the Laplace distribution for microbial co-occurrence inference. Future studies investigating sparse network constraints will need to explore the geometric implications of prior on the simplex.

This large-scale effort identified both the strengths and the limitations of the MLN applied to microbiome data. Based on classification benchmarks, we found that the MLN distribution is useful for analyzing longitudinal datasets, suggesting that these distributions could serve as a scaffold for longitudinal analysis. However, we also found that the MLN distribution performs on par with Bray-Curtis and Unifrac on cross-sectional studies. Based on our simulations, we suspect that this is due to the excessive amount of sparsity in the data, since we saw the MLN covariance begin to deteriorate at 40% sparsity, whereas the microbiome datasets benchmarked here have above 90% sparsity, even after removing features that are present in at least 30 samples. Furthermore, our training strategy is limited to the microbes that have been detected in the training dataset and future efforts will need to investigate how to account for novel microbes during prediction. We anticipate that in future extensions of this work, incorporating phylogenetic information and accounting for more confounding variables such as age and geography may help boost performance.

While we have provided some biological intuition behind the estimated co-occurrences, the biological validation of these co-occurrences is currently limited by the complexity of conducting anaerobic co-culturing experiments. Amongst the multiple co-culture studies that were investigated, few of them them had strain level taxonomies that could be detected in the full human amplicon dataset. Amplicon experiments cannot obtain strain-level resolution for most microbial taxa, but our findings suggest that strain-level resolution is critical for obtaining coherent microbe-microbe interactions. These taxonomic limitations may be resolved with strain-level shotgun metagenomics.

Finally, the many of the studies that are present in Qiita are cross-sectional studies, which will not provide a longitudinal perspective surrounding these microbial interactions. Previous studies have argued that microbe-microbe interactions can only be inferred from longitudinal data [81], citing ecological models such as Lotka-Volterra. Furthermore, previous studies have also argued that absolute abundances are necessary to infer the underlying dynamics of these microbial communities [82, 83]. We anticipate that extensions of our proposed model can integrate these sampling strategies to boost the accuracy of inferring microbe-microbe interactions.

## 5 Conclusion

Our major contribution here was providing a way to accelerate the estimation of the Multinomial Logistic Normal distribution to enable scalable infererence on large microbiome datasets. We have provided theoretical and empirical evidence to elucidate the connection between Multinomial PPCA and Multinomial VAEs. Furthermore, we have shown that this approach is competitive across multiple classification tasks, with highlighted benefits for longitudinal datasets. We anticipate that extensions of these methods will play a key role to building flexible time series models for longitudinal studies and providing accurate microbe-microbe interaction estimates for co-culture validation. We have shown-cased the merits of the proposed VAE architecture and anticipate these insights will benefit machine learning efforts across the biological sciences.

## Software Availability

Our software and pretrained models can be found at https://github.com/flatironinstitute/catvae

## Acknowledgement

We would like to thank Ian Fisk and Nick Carriero for providing the computational support required to train these models. We’d also like to acknowledge Juan Jose Egozcue and Vera Pawlowsky for discussions surrounding the ILR transform, Antonio Gonalez, Daniel McDonald and Gail Ackermann for their feedback on using Qiita and Redbiom and Adam Gayoso for contributing code to the biom repository to enable the benchmarking of LDVAE.

## A Appendix

### A.1 The ILR transform

As outlined in Equation 1, any orthogonal basis **Ψ** can be used to perform the ILR transform. Some of these bases can be represented as binary trees with *O*(*d* log *d*) elements, where the *l*th column vector of **Ψ** is given as follows:

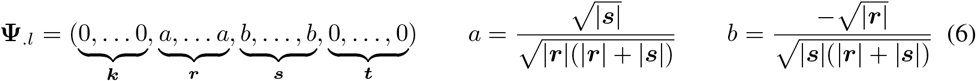

where *l* indexes an internal node in the tree with left children ***r***, right children ***s***, nodes to the left ***k*** and nodes to the right ***t*** [84] (Figure S1).

One can see that **Ψ** is a contrast matrix and can be forced to be orthonormal such that **Ψ**^***T***^ **Ψ** = ***I***_*d*_^*−*^_1_, as highlighted in Equation 6. Furthermore, this construction can be scaled to large binary trees as shown in Figure S1. Here, ***η***_*l*_ represents the log-ratios at the internal node *l*, given by

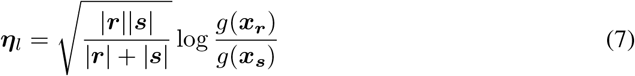

For an extended discussion how to construct contrast matrices **Ψ** see Egozcue et al [84].

Since the tree itself forms a full rank orthonormal basis and no regularization on ***W*** is used, it doesn’t matter which tree is used to parameterize the ILR basis. However, the choice of tree will influence the runtime of the ILR transform. If a balanced binary tree is used, the memory requirements representing **Ψ** can be brought down from *O*(*d*^2^) to *O*(*d* log *d*) and can reduce the matrix vector multplication runtime from *O*(*d*^2^) to *O*(*d* log *d*). This can speed up the matrix-vector multiplication operations by an order of magnitude for datasets with more than ten thousand dimensions.

The naive runtime of the ILR transform of a single sample ***x*** ∈ ℝ^*d*^ is *O*(*d*^2^) due to the running time of dense matrix-vector multiplication. As shown in Equation 1, the ILR transform can also be represented by log-linear transformation with a contrast matrix. The binary tree can be used to represent a contrast matrix, as discussed in [84].

**Figure S1:**
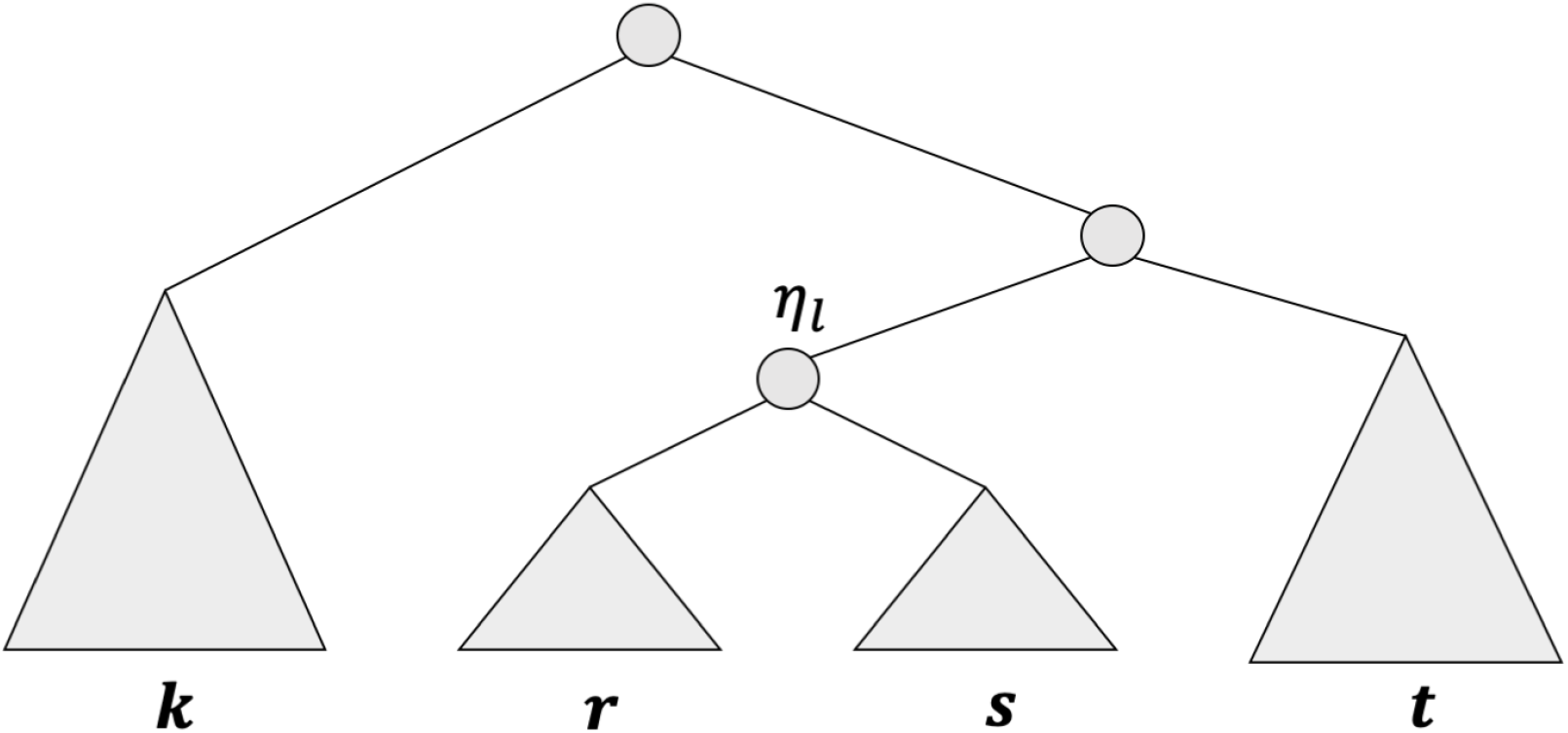
An illustration of how the ILR basis can be constructed on large trees. The quantities *g*(***x***_***r***_) and *g*(***x***_***s***_) yield the geometric means within a vector of proportion ***x*** for subsets ***x***_***r***_ and ***x***_***s***_. Here, ***r*** and ***s*** refer to the sets of features in the left and right subtrees for the internal node *l*. The log-ratios ***η*** can be obtained from either Equation 6 or Equation 7

If the binary tree is balanced, each row of **Ψ** will have *O*(log *d*) non-zero elements, since the tree has a height of *O*(log *d*). Given that there are *d* rows, the matrix-vector multiplication behind the inverse ILR transform *ϕ*(***η***) can be done in *O*(*d* log *d*). For the same reason, the ILR transform has a runtime of *O*(*d* log *d*).

### A.2 Challenges in deriving an analytical Multinomial VAE ELBO

Consider the generative model for Multinomial PPCA

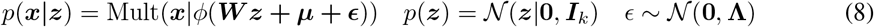

With this in mind, we wish to estimate the variational distributions *q*(***z***|***x***) to approximate the posterior *p*(***z***|***x***). This variational distribution can also be chosen to be normal distribution as follows:

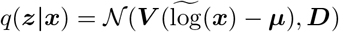

Noting that 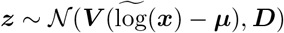, the reconstructed distribution of logits ***η* = *W*** _***z***_ **+ *μ*** is

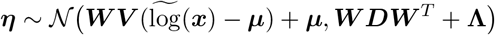

To fine-tune these variational distributions to approximate the posterior distribution, we can maximize the evidence lower bound (ELBO) given by

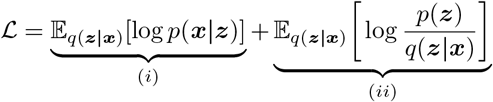

Since 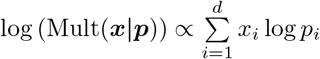, the first term (*i*) is given by

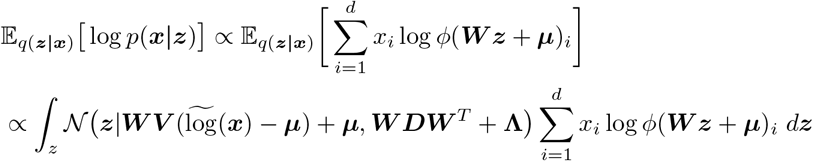

Computing the above integral is equivalent to computing the expectation of a logistic normal distribution, which does not have an analytical solution [39]. As a result, the ELBO for the Multinomial VAE is analytically intractable.

### A.3 Global Log-Concavity of the MLN Distribution

Since the MLN distribution is difficult to directly evaluate, it is challenging to make statements regarding the Multinomial VAE ELBO. However, we can make concrete statements about the posterior factors *p*(***x***|***η***) and *p*(***η***|***z***). Letting ***η*** = ***W*** _***z***_ + ***μ***, the log probability density of the multinomial distribution *p*(***x***|***η***) can be written as

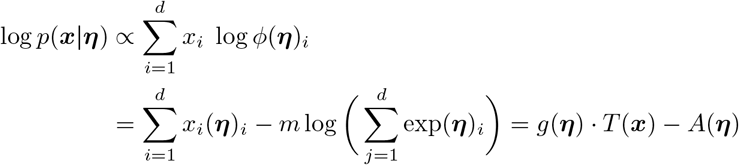

where 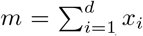 is the total number of counts and the functions *g*(***η***)_*i*_ = *ϕ*(***η***)_*i*_, *T* (***x***)_*i*_ = *x*_*i*_, and 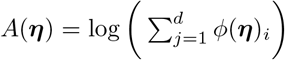 are the natural parameters of the exponential family distribution. The Hessian of log *p*(*x*|*η*) is given by

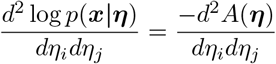

Since *A*(***η***) is strictly convex, log *p*(***x***|***η***) is strictly concave.

Similarly, the log probability density of the multivariate Gaussian distribution *p*(***η z***) = 𝒩 (***μ*, Σ**) is also strictly concave with respect to ***μ*** and **Σ**. Recall that the MLN probability density is given as

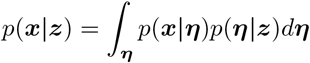

Since both *p*(***x***|***η***) and *p*(***η***|***z***) are strictly log-concave, their product is also strictly log-concave. Furthermore, since log concavity is also preserved under marginalization, thus the MLN probability density must also be strictly log-concave. Therefore, there must be a unique optimal estimate for ***μ*** and **Σ**.

### A.4 The log-concave nature of the Multinomial Linear VAE ELBO

Even though there isn’t an analyical solution to the Multinomial Linear VAE ELBO, we can still explore the properties of the ELBO. Revisiting the MLN expectation

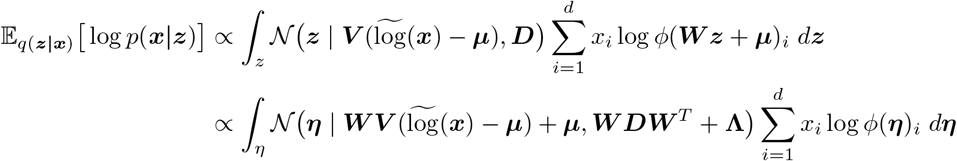

we can see that the normal distribution 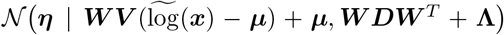 is log-concave with respect to its mean and covariance. As shown earlier, 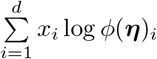 is also log-concave with respect to ***η*** = ***W*** _***z***_ + ***μ***. Since log-concavity is preserved through marginalization, 𝔼_*q*(***z***|***x***)_ log *p*(***x***|***z***) must also be log-concave with respect to the mean 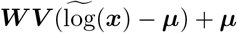 and covariance ***W DW***^*T*^ + **Λ**.

The KL divergence due to the prior

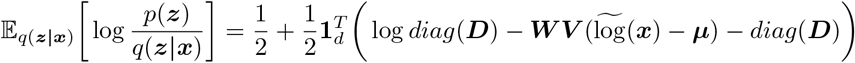

is also log-concave with respect to the mean and covariance of *q*(***z***|***x***). Since the sum of two log concave functions is also concave the ELBO is also a log concave function with respect to the mean and covariance of *q*(***z***|***x***).

Even though the Multinomial VAE ELBO itself does not offer a closed-form solution, we know that it is log-concave. This provides theoretical justification that a global optimum can be reached with reparameterization gradients. Following the intuition highlighted in [85, 24], once can see that the VAE parameters can be identified up to rotation and scale.

#### Rotation identifiability

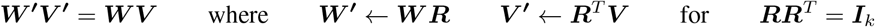

where ***R*** is any rotation matrix.

#### Scale identifiability

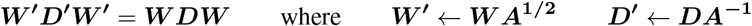

where ***D*** is a diagonal matrix with strictly positive entries.

In our VAE implementation, the eigenvalue scale identifiability issue was resolved by projecting ***W*** into a Grassmaniann manifold and fixing ***W***^***T***^ ***W*** = ***I***_*k*_ [86]. The rotation identifiability is still outstanding, and we suspect that this could explain the slow convergence rate of the Linear VAEs [49].

## B Experimental details

### B.1 Training details

All microbiome data was retrieved from Qiita using Redbiom [87]. For the mouse dataset, 39 studies were considered. Samples were excluded from the training dataset if they contained less than 1000 reads. Microbes were excluded from the entire dataset if they weren’t observed in at least 10 samples. For the human dataset, 55 studies were considered. Studies were excluded if they stored their samples at room temperature for more than 24 hours. Protocol benchmarks on DNA extraction, sample storage and the MBQC were all excluded from training. Samples were excluded from the training dataset if they contained less than 1000 reads. Microbes were excluded from the entire dataset if they weren’t observed in at least 30 samples. All samples were processed with Deblur [88] and trimmed to 100bp.

The VAEs that were trained on both the mouse and the human used very similar architectures with a 5 layer encoder with softplus activations and a dense decoder whose weighted are embedded in a Grassmannian space [86]. The latent dimensionality for both VAEs was 128 units. The only difference between the two models is the mouse VAE was trained with **Λ = *I***, whereas the human VAE was trained with a diagonal covariance **Λ** in order to better handle technical variation due to sample collection protocols. Both of these models were trained for 10k epochs, with the mouse VAE taking 6 hours and the human VAE taking 24 hours on a single Nvidia V100 GPU. Due to the long runtime, both LDVAE and LDA was only trained for 1k epochs on the mouse dataset using the default parameters in scVI and scikit-learn respectively.

The Multinomial VAEs were all trained on 80% of the full dataset. The LDVAE and LDA were also trained on the same training split for the mouse dataset.

### B.2 Co-occurrence analysis

**Figure S2:**
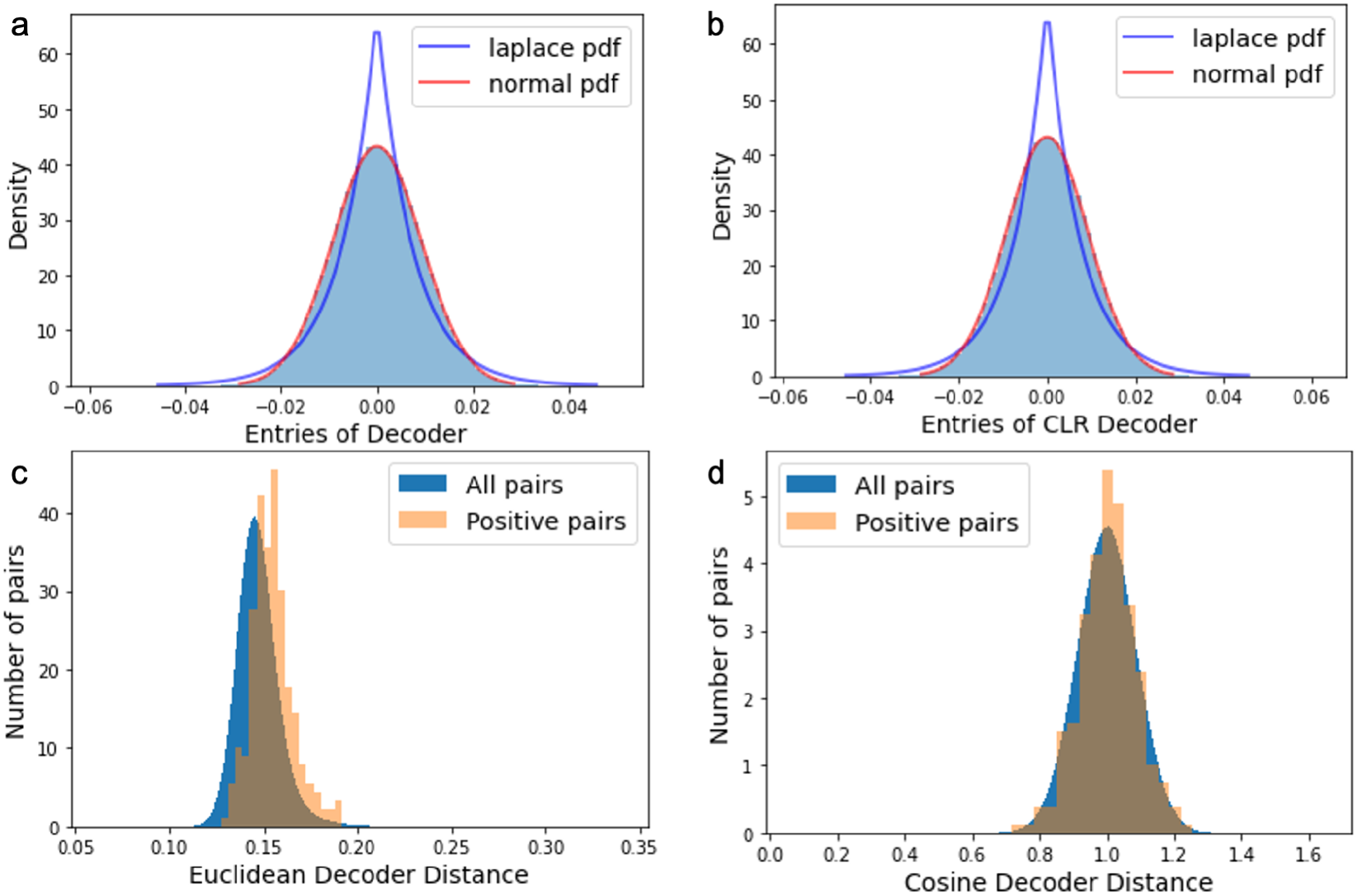
Distributions of the VAE decoder: (a) Distribution of the decoder weights in ILR space. (b) Distribution of the decoder weights after projection with **Ψ**. (c) Distribution of Euclidean distances on **Ψ*W***. (d) Distribution cosine distances from VAE decoder embedding **Ψ*W***.

